# Extensive Genetic Diversity and Host Range of Rodent-borne Coronaviruses

**DOI:** 10.1101/2020.08.11.245415

**Authors:** Wen Wang, Xian-Dan Lin, Hai-Lin Zhang, Miao-Ruo Wang, Xiao-Qing Guan, Edward C. Holmes, Yong-Zhen Zhang

## Abstract

To better understand the genetic diversity, host association and evolution of coronaviruses (CoVs) in China we analyzed a total of 696 rodents encompassing 16 different species sampled from Zhejiang and Yunnan provinces. Based on the reverse transcriptase PCR-based CoV screening CoVs of fecal samples and subsequent sequence analysis of the RdRp gene, we identified CoVs in diverse rodent species, comprising *Apodemus agrarius, Apodemus latronum, Bandicota indica, Eothenomys miletus, E. eleusis, Rattus andamanesis, Rattus norvegicus*, and *R. tanezumi. Apodemus chevrieri* was a particularly rich host, harboring 25 rodent CoVs. Genetic and phylogenetic analysis revealed the presence of three groups of CoVs carried by a range of rodents that were closely related to the Lucheng Rn rat coronavirus (LRNV), China Rattus coronavirus HKU24 (ChRCoV_HKU24) and Longquan Rl rat coronavirus (LRLV) identified previously. One newly identified *A. chevrieri*-associated virus closely related to LRNV lacked an NS2 gene. This virus had a similar genetic organization to AcCoV-JC34, recently discovered in the same rodent species in Yunnan, suggesting that it represents a new viral subtype. Notably, additional variants of LRNV were identified that contained putative nonstructural NS2b genes located downstream of the NS2 gene that were likely derived from the host genome. Recombination events were also identified in the ORF1a gene of Lijiang-71. In sum, these data reveal the substantial genetic diversity and genomic complexity of rodent-borne CoVs, and greatly extend our knowledge of these major wildlife virus reservoirs.

## 1. Introduction

Coronaviruses (CoVs) (family *Coronaviridae*, order *Nidovirales*) are important etiological agents for respiratory, enteric, hepatic, and neurological diseases of varying severity that impact a variety of animal species including humans. The first coronavirus was isolated in chicken embryos in the 1930s (Hudson and Beaudette 1932). Notably, those human coronaviruses described before 2002 were associated with mild influenza-like symptoms. However, following the emergence of SARS (Severe Acute Respiratory Syndrome) in 2002/2003, MERS (Middle East Respiratory Syndrome) in 2012 and COVID-19 (Corona Virus Disease 2019) in 2019, their potential as human pathogens has gained increasing attention. Importantly, all these diseases are associated with zoonotic CoVs, with a variety of bat species recognized as important wildlife reservoirs. In addition to bats, rodents are also a major zoonotic source of emerging viral diseases, including a number of important infectious diseases of humans (Meerburg et al. 2009; Zhang et al. 2010). Indeed, rodents are highly diverse and often live in close proximity to humans or domestic animals, presenting an increased risk after direct or indirect exposure to rodent carcasses, faces, urine and parasites.

CoVs are currently classified into four genera: *Alphacoronavirus, Betacoronavirus, Gammacoronavirus* and *Deltacoronavirus* (de Groot 2011; https://talk.ictvonline.org/taxonomy/). SARS-CoV, MERS-CoV and the agent of COVID-19, SARS-CoV-2, are all members of genus *Betacoronavirus*, itself divided into the subgenera *Embecovirus, Hibecovirus, Merbecovirus, Nobecovirus* and *Sarbecovirus*. Although rodents are important reservoirs for a range of zoonotic pathogens, prior to 2015 the only known rodent coronavirus was mouse hepatitis virus (MHV) isolated from mice in 1949 (Cheever et al. 1949). Following the discovery of four other distinct rodent coronaviruses - the alphacoronavirus Lucheng Rn rat coronavirus (LRNV) and the betacoronaviruses (subgenus *Embecovirus*) ChRCoV_HKU24, Myodes coronavirus 2JL14 (MrufCoV_2JL14) and Longquan Rl rat coronavirus (LRLV) – an increasing number of rodent associaed coronaviruses have been identified in different countries and in a range of rodent species (Wang et al. 2015; Lau et al. 2015; Tsoleridis et al. 2016; Ge et al. 2017;Wu et al. 2018). Hence, rodents are important reservoirs for members of the subgenus *Embecovirus* of betacoronaviruses and have likely played a key role in coronavirus evolution and emergence.

Zhejiang province is located in the southern part of the Yangtze River Delta on the southeast coast of China, from where rodent CoVs have previously been reported (Wang et al. 2015; Lin et al. 2017). Yunnan province is located in southern China, bordering the countries of Myanmar, Laos, and Vietnam, and is often caused the “the Kingdom of Wildlife”. A previous study from Yunnan provide identified a novel SARS-like CoV, Rs-betacoronavirus/Yunnan2013, whose ORF8 was nearly identical to ORF8 of SARS-CoVs (98% nt sequence identities) (Wu et al. 2016). Recently, two CoVs closely related to SARS-CoV-2 have been identified in *Rhinolophus* sp. (i.e. horseshoe) bats sampled from Yunnan province: RaTG13 (Zhou et al. 2020) and RmYN02 (Zhou et al. 2020). However, few rodent CoVs has been documented in Yunnan to date. To explore the diversity and characterization of CoVs in rodents, we performed a molecular evolutionary investigation of CoVs in Zhejiang province and Yunnan province, China. Our results revealed a remarkable diversity of CoVs in rodents.

## 2. Materials and Methods

### 2.1 Sample collection

This study was reviewed and approved by the ethics committee of the National Institute for Communicable Disease Control and Prevention of the Chinese CDC. All animals were kept alive after capture and treated in strictly according to the guidelines for the Laboratory Animal Use and Care from the Chinese CDC and the Rules for the Implementation of Laboratory Animal Medicine (1998) from the Ministry of Health, China, under the protocols approved by the National Institute for Communicable Disease Control and Prevention. All surgery was performed under anesthesia, and all efforts were made to minimize suffering.

All rodents were collected in 2014 and 2015 from Lijiang and Ruili cities in Yunnan province, and Longquan and Wenzhou cities in Zhejiang province, China. Sampling occurred in cages using fried food as bait set in the evening and checked the following morning. Animals were initially identified by trained field biologists, and further confirmed by sequence analysis of the mitochondrial (mt)-*cyt b* gene (Guo et al. 2013). Lung samples were collected from animals for the classification of rodent species and alimentary tract samples were collected from animals for the detection of CoVs, respectively.

### 2.2 DNA and RNA extraction

Total DNA was extracted by using the Cell & Tissue Genomic DNA Extraction Kit (Bioteke Corporation, Beijing, China) from lung samples of rodents according to the manufacturer’s protocol. Total RNA was extracted from fecal samples using the Fecal total RNA extraction kit (Bioteke Corporation, Beijing, China) according to the manufacturer’s protocol. The RNA was eluted in 50μl RNase-free water and was used as a template for further detection.

### 2.3 CoV detection and complete genome sequencing

The mt-*cyt b* gene (1140 bp) was amplified by PCR with universal primers for rodents described previously (Guo et al. 2013). CoV screening was performed using a previously published primer set by a pan-coronavirus nested PCR targeted to a conserved region of the RNA-dependent RNA polymerase gene (RdRp) gene (Wang et al. 2015). First-round reverse transcription PCR (RT-PCR) was conducted by using PrimeScript One Step RT-PCR Kit Ver.2 (TaKaRa, Dalian, China). A 10 μL reaction mixture contained 5 μL of 2 × 1 Step Buffer, 0.4 μL PrimeScript 1 Step Enzyme Mix, 0.3 μL (10μmol/l) forward primer, 0.3 μL (10μmol/l)) reverse primer, 3.5 μL RNase Free dH_2_O, and 0.5 μL of sample RNA. The PCR cycler conditions for the amplification were 50°C for 30 min (reverse transcription) then 95°C for 3 min, 35 cycle of 94°C for 45 s (denaturation), 44°C for 45 s (annealing), 72°C for 45 s (extension), then 72°C for 10 min (final extension). The PCR product was then put through a second round PCR which amplify a final PCR product of approximately 450bp.

To recover the complete viral genome, RNA was amplified by using several sets of degenerate primers designed by multiple-sequence alignments of available genome of published CoVs. Additional primers were designed according to results of the first and subsequent rounds of sequencing. The 5’ and 3’ end of the viral genome was amplified by rapid amplification of cDNA ends by using the 5’ and 3’ Smarter RACE kit (TaKaRa, Dalian, China).

RT-PCR products of expected size were subject to Sanger sequencing performed by the Sangon corporation (Beijing, China). Amplicons of more than 700 bp were sequenced in both directions. Sequences were assembled by SeqMan and manually edited to produce the final sequences of the viral genomes. Nucleotide (nt) sequence similarities and deduced amino acid (aa) similarities to GenBank database sequences were determined using BLASTn and BLASTp.

### 2.4 Phylogenetic analysis

CoV reference sequences sets representing the RdRp, S and N genes were downloaded from GenBank. Both partial RdRp gene sequences and complete amino acid sequences of the RdRp, S and N genes were used to infer phylogenetic trees. All viral sequences were aligned using the MAFFT algorithm (Katoh and Standley 2013). After alignment, gaps and ambiguously regions were removed using Gblocks (v0.91b) (Talavera and Castresana 2007). The best-fit model of nucleotide substitution was determined using jModelTest version 0.1 (Posada 2008). Phylogenetic trees were generated using the maximum likelihood (ML) method implemented in PhyML v3.0 (Guindon et al. 2010).

### 2.5 Genome recombination analysis

Potential recombination events in the history of the LRNV, LRLV and ChRCoV_HKU24 were assessed using both the RDP4 (Martin et al. 2010) and Simplot (v.3.5.1) programs. The RDP4 analysis was conducted based on the complete genome sequence, using the RDP, GENECONV, BootScan, maximum chi square, Chimera, SISCAN and 3SEQ methods within RDP4. Putative recombination events were identified with a Bonferroni corrected *P*-value cut-off of 0.01. Similarity plots were inferred using Simplot to further characterize potential recombination events, including the location of possible breakpoints.

## 3. Results

### 3.1 Collection of rodents, and the detection of CoV RNA

During 2014 and 2015 a total of 696 rodents from 16 different species were captured in Lijiang city, Ruili city, Yunnan province and Longquan city, Wenzhou city, Zhejiang province (Figure 1 and Table 1). RT-PCR was performed to detect CoVs RNA based on partial RdRp sequences. PCR products of the expected size were amplified from one *A. agrarius* collected from Longquan and two *R. norvegicus* sampled from Wenzhou; 25 *A. chevrieri*, two *A. latronum*, three *Eothenomys miletus* from Lijiang; two *B. indica*, one *E. eleusis*, one *R. andamanesis*, and two *R. tanezumi* from Ruili. Overall, 5.6% of rodents were CoV positive. All these sequences exhibited close sequence similarity to published CoVs. Specifically, two CoVs sampled from *B. indica* and one from *R. andamanensis* in Ruili shared 92.9%-96.0% nucleotide (nt) sequence similarity with LRLV; 21 CoVs from one *A. agrarius* in Longquan, one *E. eleusis* and two *R. tanezumi* in Ruili, as well as one *E. miletus*, two *A. latronum*, 12 *A. chevrieri* in Lijiang had 93.2%-98.4% nt sequence similarity to Longquan-343 and ChRCoV HKU24; 17 CoVs from two *R. norvegicus* in Wenzhou, one *A. latronum*, two *E. miletus*, and 12 *A. chevrieri* in Lijiang had 83.3%-98.9% nt sequence similarity to LRNV.

**Table 1.** Prevalence of coronaviruses in Yunnan and Zhejiang provinces China.

| Species | Yunnan |  | Zhejiang |  | Total (%) |
| --- | --- | --- | --- | --- | --- |
|  | Lijiang | Ruili | Longquan | Wenzhou |  |
| *Apodemus agrarius* | – | – | 1/44 | 0/5 | 1/49 |
| *A. chevrieri* | 25/194 | – | – | – | 25/194 |
| *A. latronum* | 2/5 | – | – | – | 2/5 |
| *Bandicota indica* | – | 2/5 | – | – | 2/5 |
| *Eothenomys miletus* | 3/119 | 0/12 | – | – | 3/131 |
| *E. eleusis* | – | 1/1 | – | – | 1/1 |
| *Microtus fortis* | – | – | 0/10 | – | 0/10 |
| *Mus musculus* | – | – | – | 0/1 | 0/1 |
| *Niviventer eha* | – | 0/2 | – | – | 0/2 |
| *N. niviventer* | – | 0/1 | 0/1 | – | 0/2 |
| *Rattus nitidus* | – | 0/2 | – | – | 0/2 |
| *R. andamanesis* | – | 1/1 | – | – | 1/1 |
| *R. losea* | – | – | 0/3 | 0/12 | 0/15 |
| *R. rattus sladeni* | – | 0/18 | – | – | 0/18 |
| *R. tanezumi* | 0/2 | 2/106 | 0/25 | 0/26 | 2/159 |
| **Total (%)** | **30/323 (9.3)** | **6/148 (4.1)** | **1/83 (1.2)** | **2/142 (1.4)** | **39/696 (5.6)** |

Note: CoV RNA positive specimens/total specimens; “–” no animals were captured.

**Figure 1.**
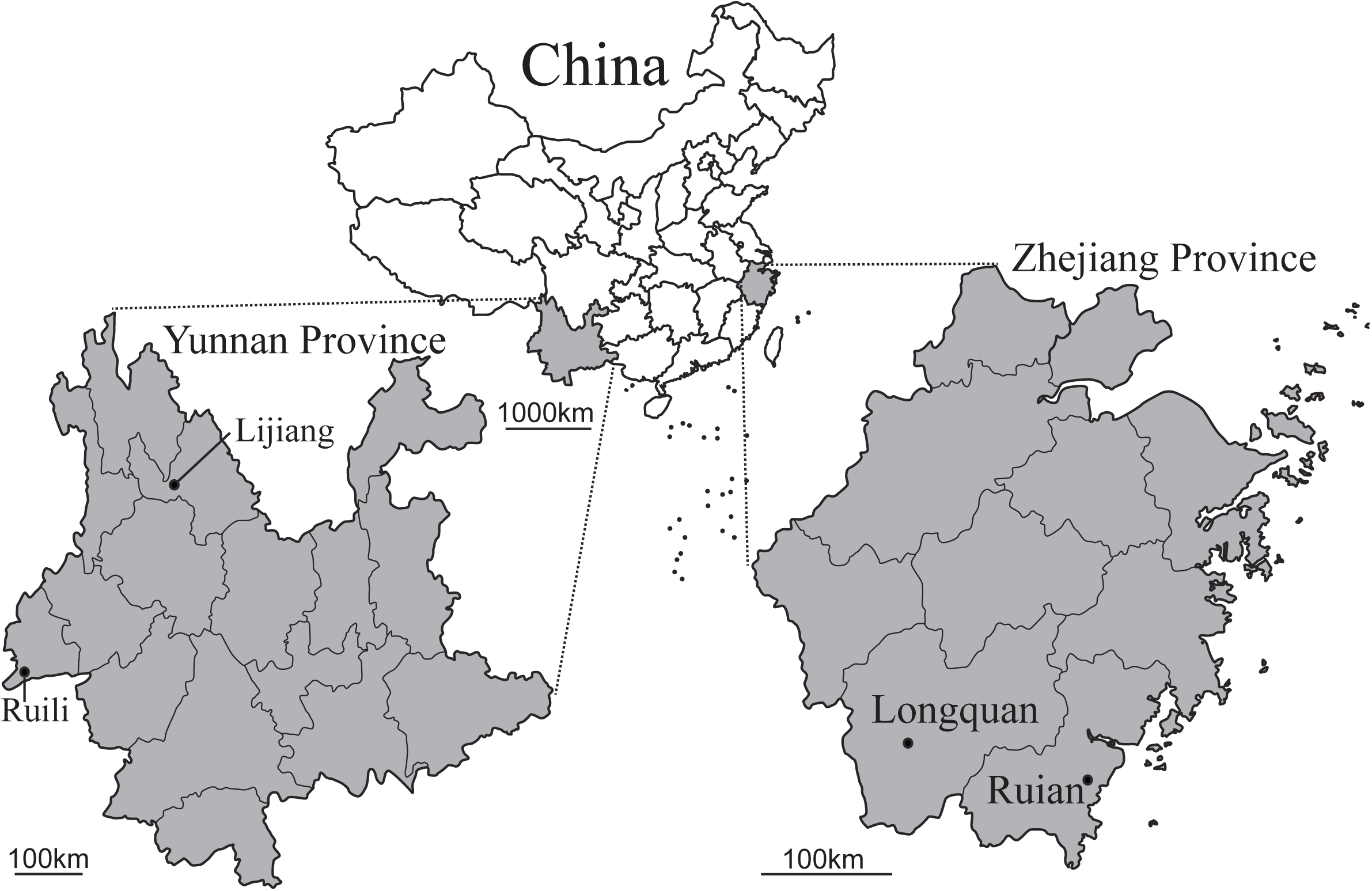
Geographic map of Yunnan and Zhejiang provinces, China, showing the location of sampling sites from where the rodents were captured.

### 3.2 Host range of rodent associated CoVs

To better understand the relationship between viruses, their hosts and their geographic distribution, we performed a phylogenetic analysis of partial RdRp (381bp) (Figure 2A). In total, 17 virus samples were identified as members of the genus *Alphacoronavirus* while 22 belonged to the genus *Betacoronavirus*. Our phylogenetic analysis revealed three different clades of rodent-borne CoVs: (i) the first clade fell within the genus *Alphacoronavirus* and contained a variety of viruses including LRNV; (ii) the second clade contained members of the subgenus *Embecovirus* (genus *Betacoronavirus*) including LRLV; (iii) the third clade also fell withing the subgenus *Embecovirus* and contained ChRCoV_HKU24. Notably, all three clades contained viruses closely related to CoVs previously identified in rodents from Zhejiang province, China.

**Figure 2.**
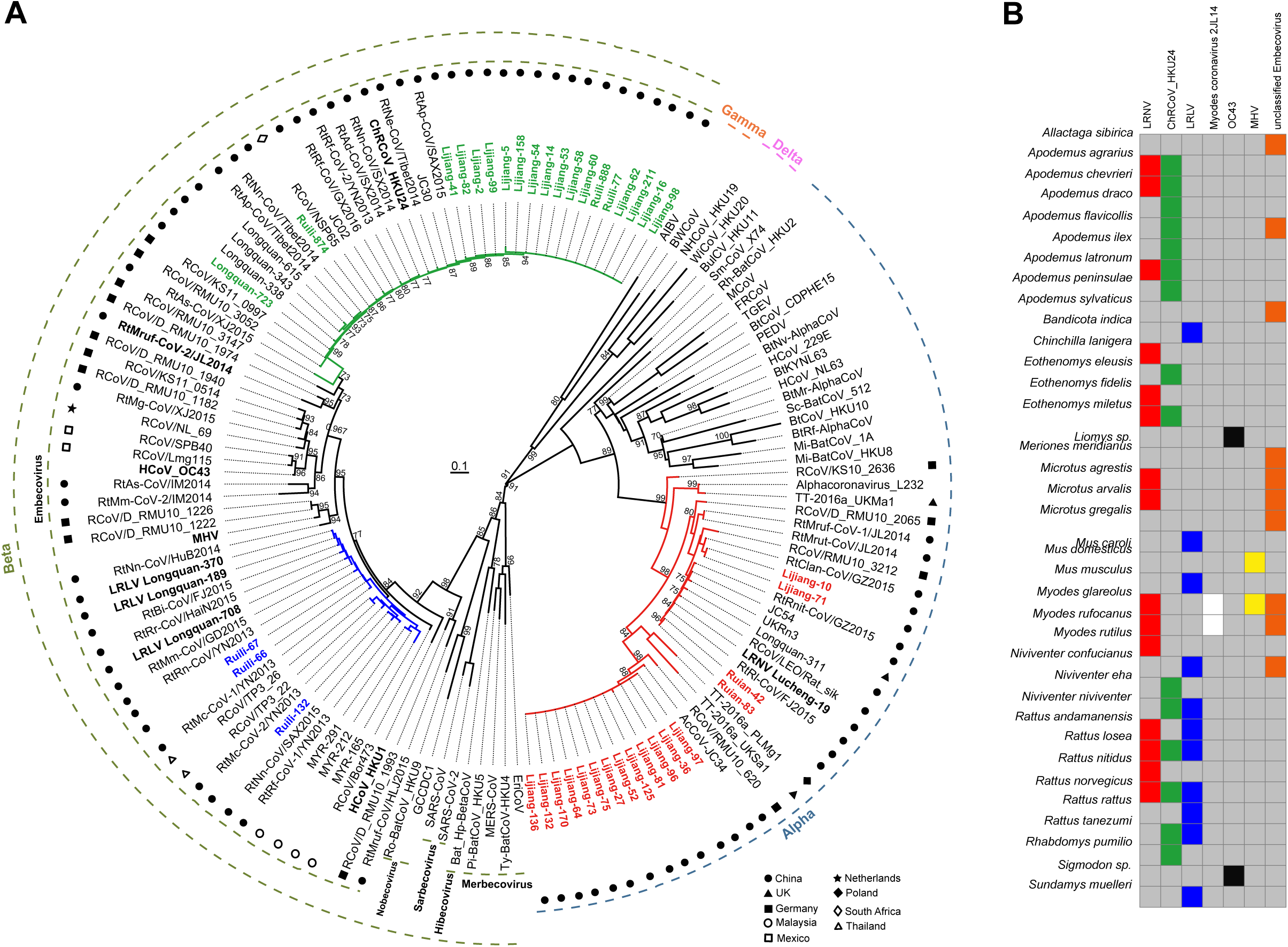
(A) Maximum likelihood phylogenetic analysis of a 381-nt sequence of the RdRp gene. Different symbols are used to indicate the country from where the viruses were identified. (B) Color block diagram of the diversity of coronaviruses carried by different rodent species.

The data generated here, along with that published previously, indicate that a total of 37 different species of rodents from nine different countries are currently known to harbor CoVs (Figure 2). Notably, every virus clade contained different rodent species, sometimes even from different subfamilies, such that there was no rigid host restriction in rodent CoVs (Figure S1). Indeed, LRNV has been identified in 15 different species in rodents, comprising *A. agrarius, A. chevrieri, A. latronum, Chinchilla lanigera, Eothenomys fidelis, E. miletus, M. agrestis, M. arvalis, M. glareolus, M. rufocanus, M. rutilus, R. lossea, R. nitidus, R. norvegicus*, and *R. sikkimensis*. In addition, multiple CoVs can be carried by the same rodent species. For example, two *Rattus* species - *R. lossea* and *R. norvegicus* - were found to carry three species of rodent-borne coronavirus. This is in contrast to previous studies in which individual CoVs were associated with a single species or genera, including *Carollia, Eptesicus, Miniopterus, Scotophilus*, and *Rhinolipus* bats (Anthony et al.2013; Drexler et al.2010; Fischer et al.2016; Wacharapluesadee et al. 2015).

### 3.3 Characterization of viral genomes

To better characterize the CoVs found in this study, complete or nearly complete genome sequences data for the three variants of LRNV (Lijiang-71, Lijiang-170, Wenzhou-83) and four variants of ChRCoV_HKU24 (Lijiang-41, Lijiang-53, Ruili-874, Longquan-723) as well as the single variant of LRLV (Ruili-66) were obtained by assembly of the sequences of the RT-PCR products from the RNA directly extracted from the individual specimens.

The three genomes of LRNV shared 77.5%-92.4% nt sequence similarity with each other. The genome sizes of Wenzhou-83, Lijiang-71 and Lijiang-170 were 28599, 29147 and 27563 nt, respectively, with the G+C contents of 40.29%, 39.35% and 40.21%. The genomes of Wenzhou-83, Lijiang-71, Lijiang-170 had 97.7%, 92.3% and 77.6% overall nucleotide identity with LRNV Lucheng-19, respectively. The genome organization was similar to that of other LRNVs and had the characteristic gene order 5’-replicase ORF1ab, spike (S), envelope (E), membrane (M), nucleocapsid (N)-3’. Strikingly, however, a major difference among these LRNVs is the additional ORF(s) encoding nonstructural (NS) proteins and located between ORF1ab and S gene (Figure 3). According to the presence and quantity of non-structural proteins, the virus can be divided into three genomic variants: (i) the first variant comprised Lijiang-170 and AcCoV-JC34 in which no NS protein was observed; (ii) the second variant contains Ruian-83, Lucheng-19 and RtRl-CoV/FJ2015 for which there is a putative NS2 gene between their ORF1ab and S gene. The putative NS2 gene lengths for Ruian-83, Lucheng-19 and RtRl-CoV/FJ2015 were 825, 828 and 825 nt, respectively, with 93.5-94.8% sequence identity; (iii) the third variant comprises Lijiang-71, RtClan-CoV/GZ2015 and RtMruf-CoV-1/JL2014 and contains two putative non-structural proteins - NS2 and NS2b – located between the ORF1ab and S gene. The putative NS2 gene of the third variant is 828 nt in length, with 82.9-88.9% sequence identity. Similarly, the putative NS2b has a gene length of 462 nt and exhibits 77.5-96.3% sequence identity among these three viruses. Strikingly, a blastp search reveals that the NS2b encodes a putative nonstructural protein of 153 amino acid residues in length that has no amino acid sequence similarity to other coronaviruses; rather, this sequence exhibits ∼43% amino acid identity to the C-type lectin-like protein within the rodent *Microtus ochrogaster* genome. Hence, this pattern suggests that the NS2b gene may have originally been acquired from the host genome during evolutionary history. Moreover, the amino acid sequence identity between Lijiang-170, Lijiang-71, Ruian-83 and LRNV was greater than 90% in RdRp, E, and M genes (as expected from members of the same species), but only 70%-88% in ADRP, 3CLpro, ORF1ab and S (Table 2). Further analysis of the characteristics of Lijiang-170, for which a complete genome sequence is available, shows that it has similar transcription regulatory sequence (TRS) to AcCoV-JC34 (Table 3). Hence, Lijiang-170 and AcCoV-JC34 may represent a novel subtype of LRNV that exhibits marked differences to the prototype strain Lucheng-19.

**Table 2.** Percent amino acid sequence identity between Lijiang-170 and known alpha-CoVs.

|  | ADRP | 3CL<sup>pro</sup> | RdRp | Hel | ORF1ab | S | E | M | N |
| --- | --- | --- | --- | --- | --- | --- | --- | --- | --- |
| **Lijiang-170** |  |  |  |  |  |  |  |  |  |
| Lijiang-71 | 74.7 | 88.3 | 94.4 | 95.8 | 85.0 | 69.1 | 93.6 | 92.7 | 80.5 |
| Ruian-83 | 71.3 | 88.3 | 94.6 | 96.5 | 85.0 | 69.5 | 93.6 | 92.3 | 78.5 |
| LRNV Lucheng-19 | 72.4 | 88.0 | 94.7 | 96.3 | 85.1 | 71.0 | 93.6 | 92.3 | 78.1 |
| Rh-BatCoV HKU2 | 44.8 | 59.0 | 74.3 | 72.9 | 53.4 | 41.3 | 36.0 | 56.6 | 30.3 |
| HCoV-229E | 42.5 | 54.7 | 73.0 | 74.2 | 51.6 | 21.0 | 37.3 | 50.0 | 29.2 |
| HCoV-NL63 | 43.7 | 55.7 | 73.3 | 72.7 | 51.7 | 21.4 | 36.0 | 53.8 | 30.3 |
| Mi-BatCoV 1A | 46.0 | 55.0 | 76.2 | 73.5 | 51.3 | 22.1 | 36.5 | 52.3 | 28.5 |
| Ro-BatCoV HKU10 | 43.7 | 55.7 | 74.8 | 73.5 | 52.1 | 23.4 | 32.0 | 57.5 | 31.3 |
| PEDV | 42.5 | 56.0 | 74.1 | 71.7 | 52.6 | 23.0 | 35.1 | 56.9 | 31.0 |
| Sc-BatCoV 512 | 47.1 | 55.3 | 73.0 | 71.7 | 52.3 | 23.4 | 36.5 | 56.6 | 31.0 |
| TGEV | 36.8 | 56.6 | 76.2 | 72.0 | 52.1 | 22.0 | 38.5 | 56.9 | 28.7 |
| MCoV | 34.5 | 59.3 | 77.4 | 72.5 | 52.1 | 22.1 | 42.3 | 52.0 | 28.5 |
| FCoV | 37.9 | 57.7 | 75.1 | 71.5 | 51.5 | 21.5 | 43.6 | 56.1 | 28.7 |
| CCoV | 35.6 | 58.7 | 76.1 | 72.0 | 51.9 | 22.5 | 42.3 | 57.3 | 30.3 |

**Table 3.**
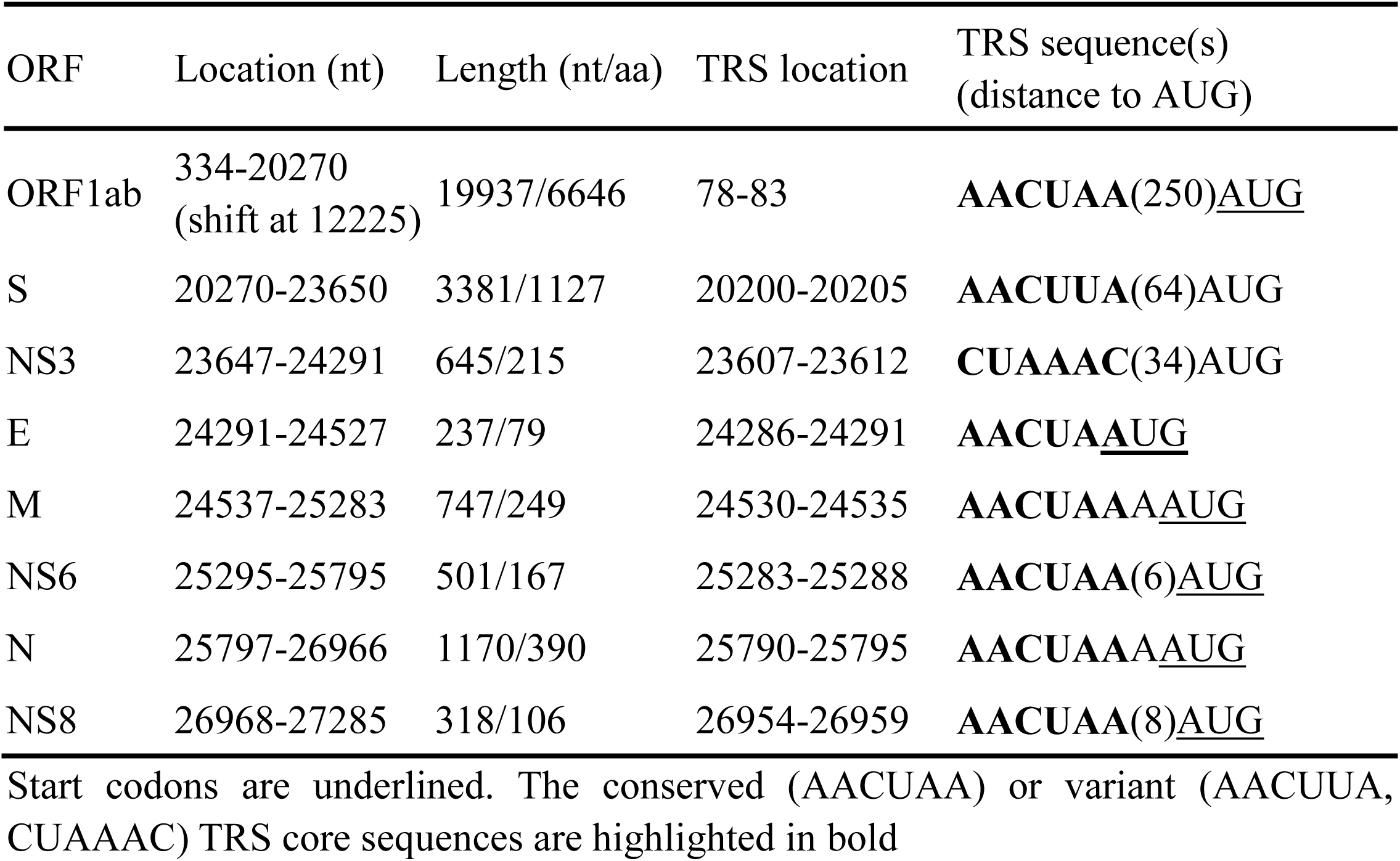
Locations of predicted ORFs in the genome of Lijiang-170.

**Figure 3.**
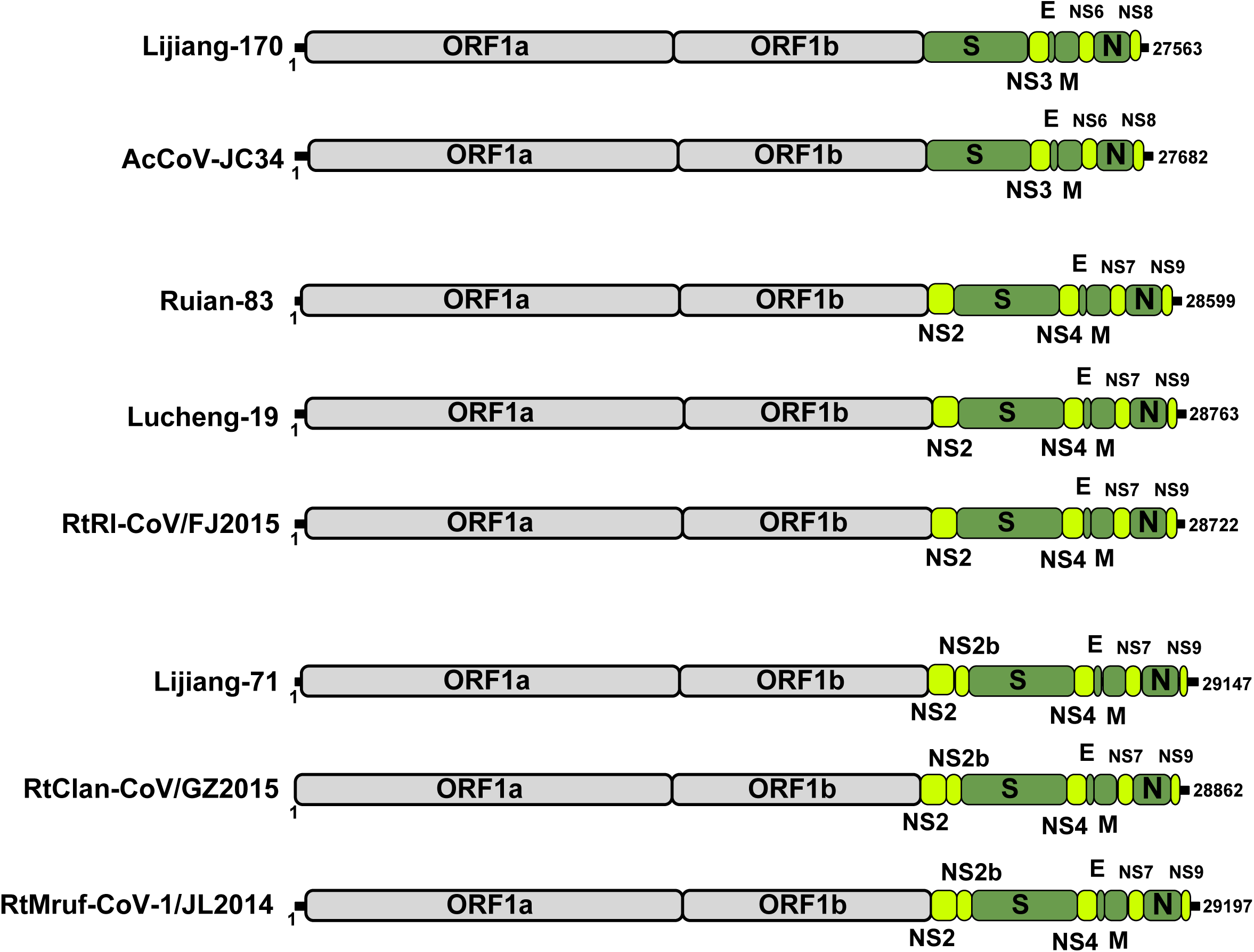
Comparison of genome organizations of the different coronaviruses identified here. All genomes are drawn to scale.

In contrast, Lijiang-53, Lijiang-41, Ruili-874, Longquan-723 were most closely related to ChRCoV_HKU24, exhibiting 94.0%-96.1% nt sequence similarity. Strikingly, the length of nsp3 in Lijiang-41 differed from those of ChRCoV_HKU24 as a result of a 75 nt deletion. Ruili-66 was most closed to LRLV and shared 92.7% nt sequence similarity with Longquan-370 and Longquan-189 of LRLV found in Longquan.

### 3.4 Phylogenetic analysis of viral sequences

To better understand the evolutionary relationships among the CoVs described here and those identified previously, we estimated phylogenetic trees based on the amino acid sequences of the RdRp, S and N proteins. The analysis of all three proteins from the LRNV clade again suggests that LRNV can be divided into two phylogenetic subtypes (I and II); indicated on Figure 4). Indeed, there was a clear division phylogenetic between the subtype I and II LRNV sequences in the RdRp, S and N amino acid trees, and while intra-subtype (I or II) sequences shared high nucleotide sequence identities (92.4% - 97.7%), inter-subtype sequence identity was only ∼77.5%. Notably, this phylogenetic analysis also suggested that clade LRNV had a recombinant evolutionary history: while the LRNV clade formed a distinct lineage in the RdRp and N gene trees (although with little phylogenetic resolution in the latter), it clustered with *Rhinolophus chevieri* coronavirus HKU2, BtRf-AlphaCoV/YN2012 and Sm-CoV X74 in the S gene tree. In contrast, the clade 2 and clade 3 rodent CoVs were consistently closely related to LRLV and ChRCoV_HKU24 in the RdRp, S and N amino acid trees.

**Figure 4.**
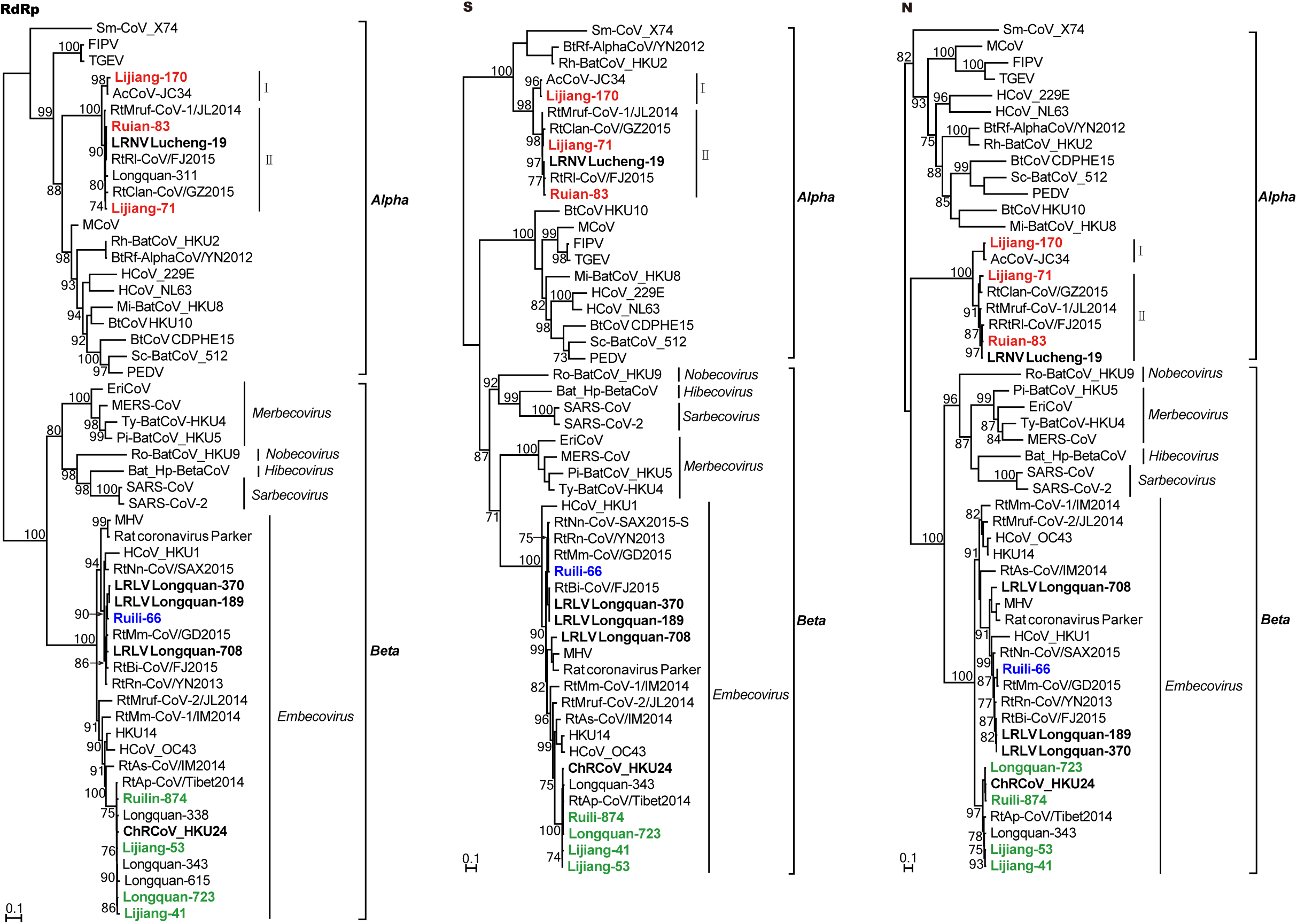
Maximum likelihood phylogenetic analysis of the RdRp, S and N proteins of Lijiang-41, Lijiang-53, Lijiang-71, Lijiang-170, Wenzhou-83, Longquan-723, Ruili-66, and Ruili-874. Numbers above individual branches indicate the percentage bootstrap support (1000 replicates). For clarity, bootstrap support values are shown for key internal nodes only. The scale bar represents the number of amino acid substitutions per site. The trees were rooted between the alpha- and beta-CoVs.

### 3.5 Recombination

Multiple methods within the RDP program (Martin et al. 2010) identified statistically significant recombination events in Lijiang-71 (*p* <3.05×10 ^-23^ to *p* <7.11×10^−13^) (Figure 5). When Lijiang-71 was used as the query for sliding window analysis with RtClan-CoV/GZ2015 and Lucheng-19 as potential parental sequences, four recombination breakpoints at nucleotide positions 8,188, 8,636, 9,030 and 11,251 in the sequence alignment were observed. This pattern of recombination events is further supported by phylogenetic and similarity plot analyses (Figure 5). Specifically, in the major parental region (1-8,187, 8,637-9,029 and 11,252-29,349), Lijiang-71was most closely related to RtClan-CoV/GZ2015, while in the minor parental region (8188-8636 and 9,030-11,251) it was more closely related to Lucheng-19.

**Figure 5.**
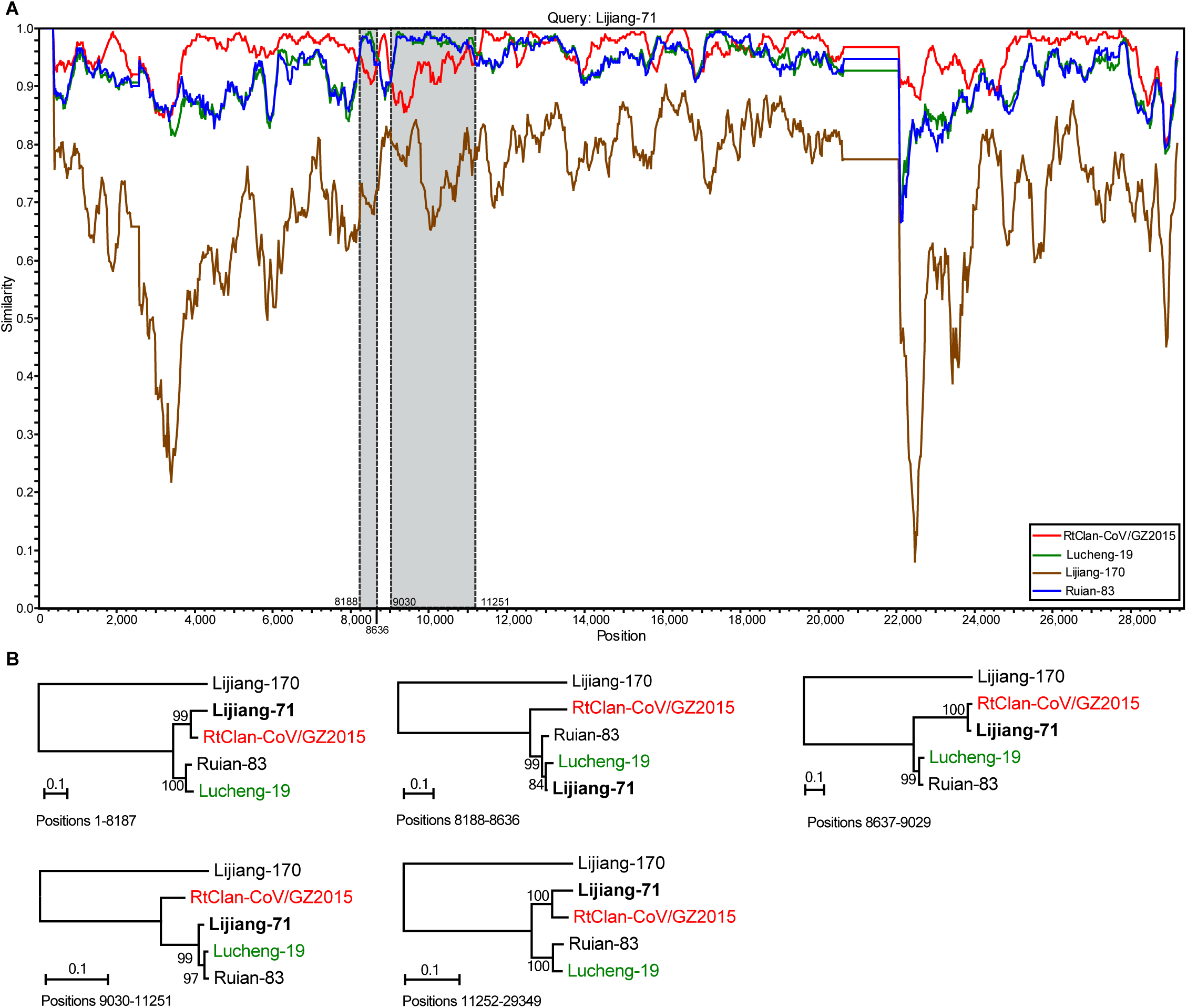
Putative recombination events within the genome of Lijiang-71. (a) The sequence similarity plot reveals four putative recombination breakpoints (black dashed lines), with their locations indicated at the bottom. The plot shows genome scale similarity comparisons of the Lijiang-71 (query) against RtClan-CoV/GZ2015 (major parent, red) and Lucheng-19 (minor parent, green). The background color of major parental region is white, while that of minor parental region is gray. (b) Phylogenies of the major parental region (positions 1-8,187, 8,637-9,029 and 11,252-29,349) and minor parental region (positions 8188-8636 and 9030-11251). Phylogenies were estimated using a maximum likelihood method and were mid-point rooted for clarity only. Numbers above or below the branches indicate percentage bootstrap values. The scale bar represents the number of substitutions per site.

## 4. Discussion

We screened CoVs in 696 rodents from 16 different species sampled at four sites in Zhejiang and Yunnan provinces, China. Overall positivity rates were approximately 6%, although they ranged from 10.2% in Lijiang city to only 1.3% in Zhejiang province. The latter is lower than the CoV detection rates described in a previous study undertaken in Zhejiang province despite the use of similar methodologies (Wang et al., 2015). We found that *A. chevrieri* had a relatively high CoV detection rate (25/194, 12.89%) in Lijiang city, Yunnan province, consistent with a previous study showing that *A. chevrieri* had a high detection rate of CoV (21/98, 21.4%) in Jianchuan county, also in Yunnan province (Ge et al. 2017). As *A. chevrieri* is a dominant species in Lijiang city, such a high coronavirus infection rate highlights the need for ongoing surveillance.

Our analysis of rodent-borne CoVs revealed that all currently recognized viruses fall into six groups - LRNV, LRLV, ChRCoV_HKU24, Myodes coronavirus 2JL14, HCoV_OC43, MHV and unclassified members of the genus *Embecovirus*. Phylogenetic analysis based on the partial RdRp gene reveals that coronaviruses from different countries (China, UK, Germany, Malaysia, Mexico, Netherland, Poland, South Africa and Thailand) group together with no obvious geographic pattern. Indeed, at least 37 different rodent species are known to carry coronaviruses, such that they have an extensive host range in these animals, and with frequent cross-species transmission. A growing number of rodent species are found to carry members of the genus *Embecovirus*, indicating that rodents indeed played an important role for *Embecovirus* spread and evolution, and are the likely reservoir hosts for the human coronavirus HKU1 (Woo et al. 2005).

Our analysis also provided evidence for multiple variants of rodent-borne CoVs in the genus *Alphacoronavirus* that different in the genome organization. For example, although Lijiang-170 had greatest sequence similarity to LRNV variant Lucheng-19, it does not contain a NS2 between ORF 1b and S gene as observed in most other alphacoronaviruses. More striking was that three viruses (Lijiang-71, RtClan-CoV/GZ2015 and RtMruf-CoV-1/JL2014) had a putative nonstructural protein of 153 amino acids located downstream of NS2 that likely resulted from a past horizontal gene transfer event involving the rodent host genome. In addition, inter-virus recombination events were identified in Lijiang-71. Such mechanisms of genetic transfer may ultimately lead to the creation of novel viruses, perhaps with variable phenotypic properties (Su et al. 2016).

In conclusion, our study revealed a high diversity of CoVs circulating in rodents from Yunnan and Zhejiang provinces, China, including the discovery of a putative novel viral subtype and new rodent host species. Undoubtedly, the larger scale surveillance and analyses of CoV infections in rodents is required to better understand their genetic diversity, cellular receptors, inter-host transmission and evolutionary history.

## Data availability

The eight complete or nearly complete CoVs genome sequences generated in this study have been deposited in the GenBank database under the accession numbers MT820625-MT820632.

## Figure legends

**Figure S1**. Tanglegram depicting the evolutionary associations between rodent associated CoVs and their hosts. The virus tree was estimated using the RdRp gene (right) and the host tree (left) was based on topology implied in the Time tree of life (http://www.timetree.org/).

