## Supplementary figures and images for "Extensive Genetic Diversity and Host Range of Rodent-borne Coronaviruses"

### Supplemental Figure S1

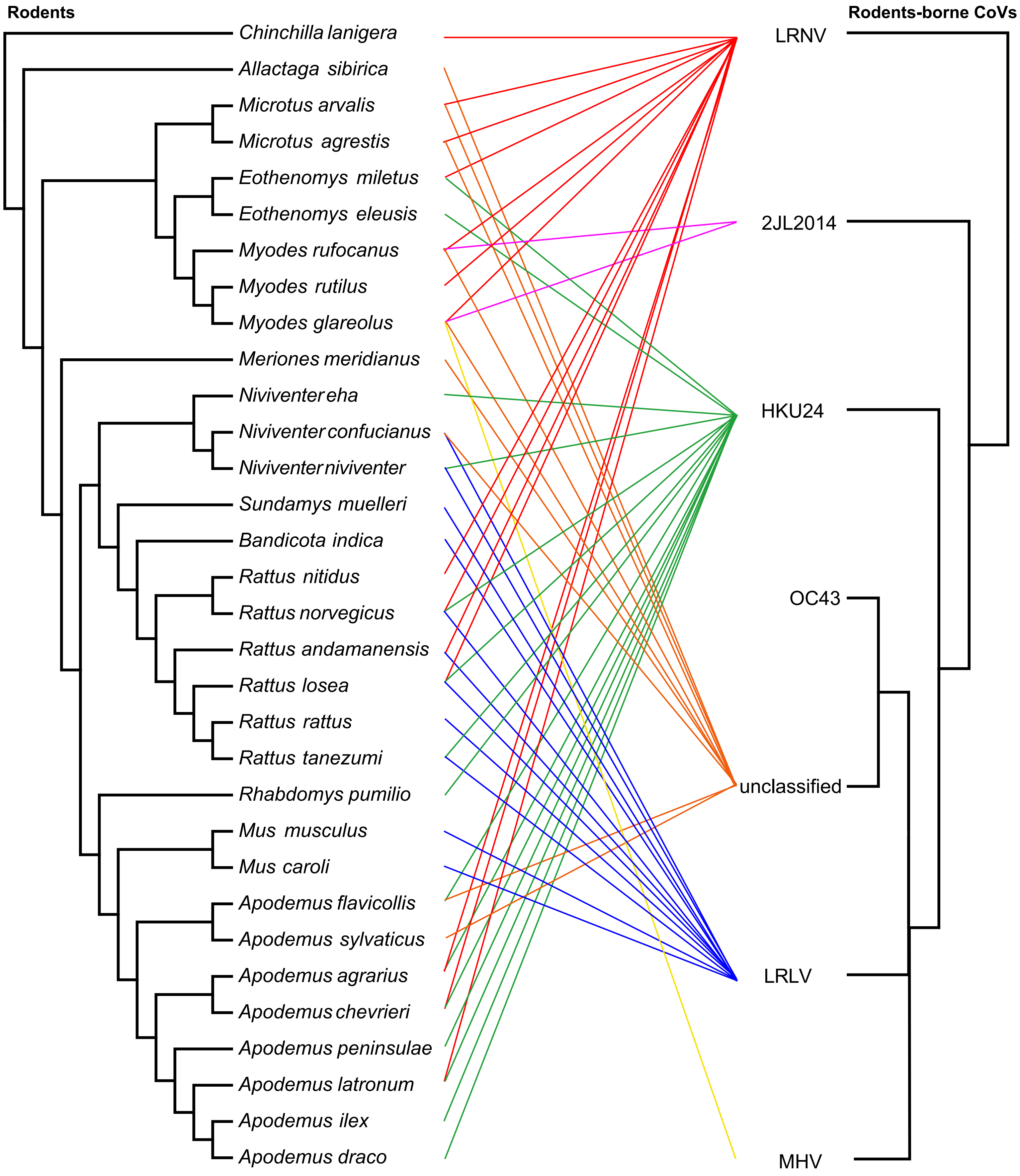
